# TCRseek: Scalable Approximate Nearest Neighbor Search for T-Cell Receptor Repertoires via Windowed k-mer Embeddings

**DOI:** 10.64898/2026.03.20.713313

**Authors:** Yang Yang

## Abstract

The rapid growth of T-cell receptor (TCR) sequencing data has created an urgent need for computational methods that can efficiently search CDR3 sequences at scale. Existing approaches either rely on exact pairwise distance computation, which scales quadratically with repertoire size, or employ heuristic grouping that sacrifices sensitivity. Here we present TCRseek, a two-stage retrieval framework that combines biologically informed sequence embeddings with approximate nearest neighbor (ANN) indexing for scalable search over TCR repertoires. TCRseek first encodes CDR3 amino acid sequences into fixed-length numerical vectors through a multi-scale windowed k-mer embedding scheme derived from BLOSUM62 eigendecomposition, then indexes these vectors using FAISS-based structures (IVF-Flat, IVF-PQ, or HNSW-Flat) that support sublinear-time search. A second-stage reranking module refines the shortlisted candidates using exact sequence alignment scores (Needleman–Wunsch with BLOSUM62), Levenshtein distance, or Hamming distance. We benchmarked TCRseek against tcrdist3, TCRMatch, and GIANA on a 100,000-sequence corpus with precomputed exact ground truth under three distance metrics. Under cross-metric evaluation—where the reranking and ground truth metrics differ, providing the most informative test of generalization—TCRseek achieved NDCG@10 = 0.890 (Levenshtein ground truth) and 0.880 (Hamming ground truth), ranking highest among the retained baselines under Hamming and remaining competitive with tcrdist3 (0.894) under Levenshtein. When the reranking metric matches the ground truth definition (BLOSUM62 alignment), NDCG@10 reached 0.993, confirming that the ANN shortlist captures *>*99% of true neighbors—the expected ceiling of the two-stage design. On the 100,000-sequence corpus, TCRseek achieved 3.6–39.6× speedup over exact brute-force search depending on index type and distance metric, with the largest gains for alignment-based retrieval. These results demonstrate that embedding-based ANN search provides a practical and scalable alternative for TCR repertoire analysis.

## 1 Introduction

Adaptive immunity relies on the extraordinary diversity of T-cell receptors, whose complementarity-determining region 3 (CDR3) loops constitute the primary determinant of antigen recognition specificity. High-throughput TCR sequencing technologies now routinely generate datasets comprising millions of unique CDR3 sequences from individual patients, enabling population-scale studies of immune responses to infections, vaccines, autoimmune disorders, and cancer immunotherapy (Robins et al., 2009; Emerson et al., 2017; DeWitt et al., 2018). A central computational challenge in analyzing these data is the identification of TCR sequences that share functional similarity—sequences recognizing the same peptide-major histocompatibility complex (pMHC) target—within and across repertoires. Solving this problem at scale is essential for applications ranging from epitope deconvolution and vaccine monitoring to biomarker discovery and adoptive T-cell therapy design.

Several computational methods have been developed to quantify TCR sequence similarity and identify groups of functionally related receptors. TCRdist, introduced by Dash et al. (2017), defines a biologically motivated distance metric based on position-weighted BLOSUM62 alignment scores across all CDR loops; its successor tcrdist3 (Mayer-Blackwell et al., 2021) provides efficient sparse-distance computation for moderate-scale analyses. ClusTCR (Valkiers et al., 2021) groups CDR3 sequences by length and Hamming distance before applying Markov clustering, achieving fast runtime on moderately sized datasets. GLIPH and GLIPH2 (Glanville et al., 2017; Huang et al., 2020) identify conserved motifs shared among TCR sequences to infer antigen specificity groups. GIANA (Zhang et al., 2021) uses isometric encoding to transform CDR3 sequences into numerical representations for rapid clustering. TCRMatch (Chronister et al., 2021) employs a scoring matrix approach for database-scale epitope assignment. More recently, deep learning methods such as DeepTCR (Sidhom et al., 2021), TCR-BERT (Wu et al., 2021), and TouCAN (Drost et al., 2024) have demonstrated the potential of learned representations for TCR specificity prediction.

Despite these advances, a fundamental scalability bottleneck remains. Methods based on exact pairwise distance computation—whether using edit distances, alignment scores, or learned embeddings—exhibit *O*(*N* ^2^) time complexity for all-against-all comparison over *N* sequences, rendering them impractical for modern repertoire datasets that may contain 10^6^ to 10^8^ unique CDR3 sequences. Even approaches that restrict comparison to same-length sequences (e.g., Hamming-based methods) face quadratic scaling within length groups. Motif-based methods such as GLIPH2 avoid this scaling problem but sacrifice the ability to provide ranked similarity lists and continuous distance measures. The field therefore lacks a method that simultaneously offers ranked nearest-neighbor retrieval, biologically meaningful distance quantification, and sublinear query-time scaling.

Approximate nearest neighbor (ANN) search, a mature field in computer science and information retrieval, offers a principled solution to this scalability challenge. Locality-sensitive hashing (LSH) and related indexing structures—including inverted file systems (IVF), product quantization (PQ), and hierarchical navigable small world (HNSW) graphs—enable search over millions of high-dimensional vectors with query times that are logarithmic or sublinear in database size, at the cost of a bounded approximation error (Jegou et al., 2011; Malkov & Yashunin, 2020). The FAISS library (Johnson et al., 2019) provides optimized GPU and CPU implementations of these structures, supporting billion-scale vector search in production systems. However, applying ANN search to TCR sequences requires an embedding function that maps variable-length amino acid sequences into fixed-length numerical vectors such that proximity in the embedding space reflects biological similarity—a requirement that standard k-mer counting or one-hot encoding approaches do not adequately satisfy.

In this work, we introduce TCRseek, a two-stage framework for scalable TCR repertoire retrieval that bridges this gap (Fig. 1). The first stage transforms CDR3 sequences into fixed-length vectors through a multi-scale windowed k-mer embedding that captures both local amino acid composition and positional information along the CDR3 loop. Amino acid residues are represented using vectors derived from BLOSUM62 eigendecomposition, preserving physicochemical substitution patterns that are directly relevant to TCR-pMHC recognition. K-mers of multiple sizes (*k* = 3, 4, 5) are assigned to positional windows (*B* = 3, 5, 10) along the sequence, and the resulting aggregate vectors are concatenated and L2-normalized to produce a single embedding per CDR3. These embeddings are then indexed using FAISS structures (IVF-Flat, IVF-PQ, or HNSW-Flat) for sublinear-time approximate nearest neighbor search. The second stage applies exact reranking of the ANN shortlist using biologically rigorous scoring functions— Needleman–Wunsch global alignment with BLOSUM62 substitution matrix, Levenshtein edit distance, or Hamming distance—to ensure that the final output reflects true sequence-level similarity rather than embedding-space artifacts.

**Figure 1:**
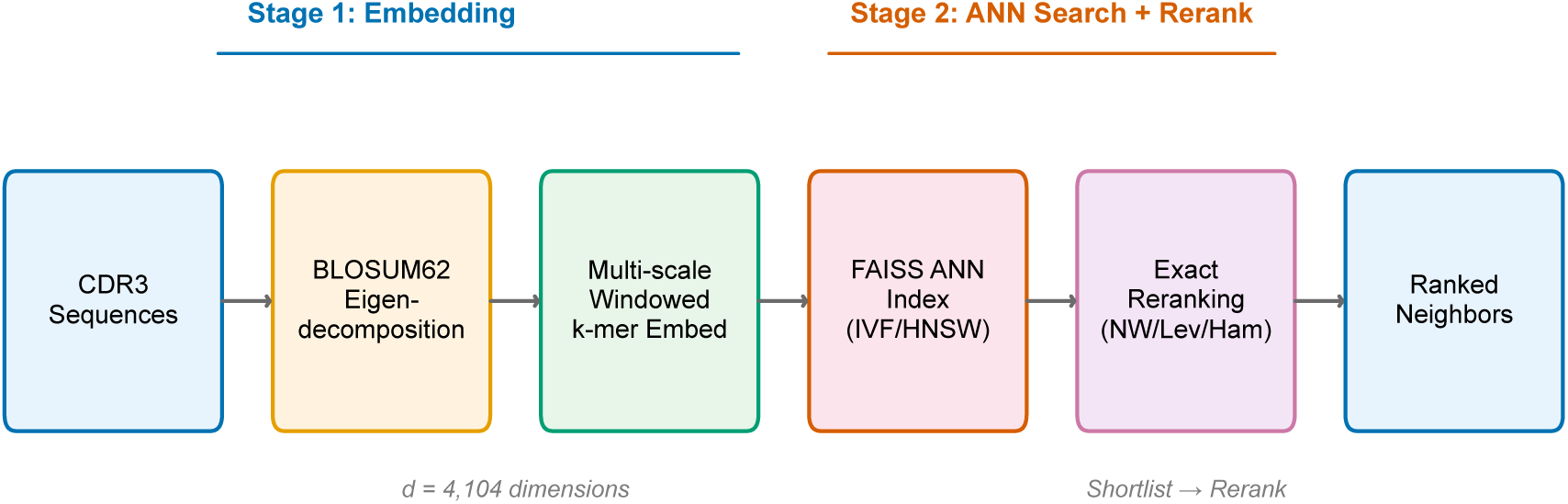
TCRseek two-stage retrieval pipeline. CDR3 amino acid sequences are first encoded into 4,104-dimensional numerical vectors through BLOSUM62 eigendecomposition and multi-scale windowed k-mer embedding (Stage 1), then indexed using FAISS approximate nearest neighbor structures. Retrieved candidates are reranked using exact sequence-level scoring—Needleman–Wunsch alignment, Levenshtein distance, or Hamming distance—to produce a final ranked neighbor list (Stage 2).

We evaluated TCRseek through a comprehensive benchmark framework that compares retrieval accuracy, computational efficiency, and scalability against existing TCR analysis tools including tcrdist3 (Mayer-Blackwell et al., 2021), TCRMatch (Chronister et al., 2021), and GIANA (Zhang et al., 2021). In a head-to-head multi-method comparison on 100,000 CDR3 sequences with precomputed exact ground truth under three distance metrics, TCRseek achieved the strongest matched-metric retrieval performance and competitive cross-metric generalization, with 3.6–39.6× speedup over exact brute-force search. Our results establish that ANN-based retrieval, when combined with biologically grounded embeddings and exact reranking, provides a practical path toward repertoire-scale TCR analysis.

## 2 Results

### 2.1 Embedding Preserves BLOSUM62 Substitution Patterns

We first validated that the BLOSUM62 eigendecomposition produces amino acid vectors whose dot-product similarities faithfully reconstruct the original substitution matrix. Retaining *d* = 19 positive eigenvalues and discarding the single numerical zero mode of the centered 20 × 20 BLOSUM62 matrix yielded a reconstruction with off-diagonal Pearson correlation *>* 0.99 and RMSE *<* 0.1, confirming that no meaningful substitution information is lost in the embedding.

### 2.2 Benchmark Setup and Ground Truth

We evaluated TCRseek on a ground-truth-locked benchmark derived from a large-scale TCR sequencing dataset comprising 10,000,000 total input rows and 7,289,308 unique valid CDR3 sequences. A deterministic hash-based sampling procedure selected 100,000 unique sequences as the benchmark corpus. Exact top-200 nearest neighbors were precomputed for every sequence under three distance metrics—Hamming-like distance, Levenshtein edit distance, and affine-gap BLOSUM62 global alignment distance—requiring 17,776.61 seconds of computation.

Three FAISS index configurations were evaluated: IVF-Flat (nlist = 256, nprobe ∈ {1, 4, 8, 16}), IVF-PQ (nlist = 256, *m* = 20, nbits = 8, nprobe ∈ {1, 4, 8, 16}), and HNSW-Flat (*M* = 32, efConstruction = 200, efSearch ∈ {32, 64, 128}).

### 2.3 ANN-Only Retrieval Performance

We first assessed the retrieval quality of the embedding and ANN index alone, without reranking (Table 1, Fig. 2). HNSW-Flat achieved the highest ANN-only recall across all three metrics: 0.3865 (Hamming), 0.3636 (Levenshtein), and 0.5900 (alignment). IVF-Flat achieved comparable recall at 0.3749, 0.3588, and 0.5746, respectively. IVF-PQ showed lower recall (0.3018, 0.2790, 0.4618) due to the lossy nature of product quantization, but offered dramatically lower query latency at 0.0324 ms median per query compared to 0.9324 ms (IVF-Flat) and 0.7382 ms (HNSW-Flat).

**Figure 2:**
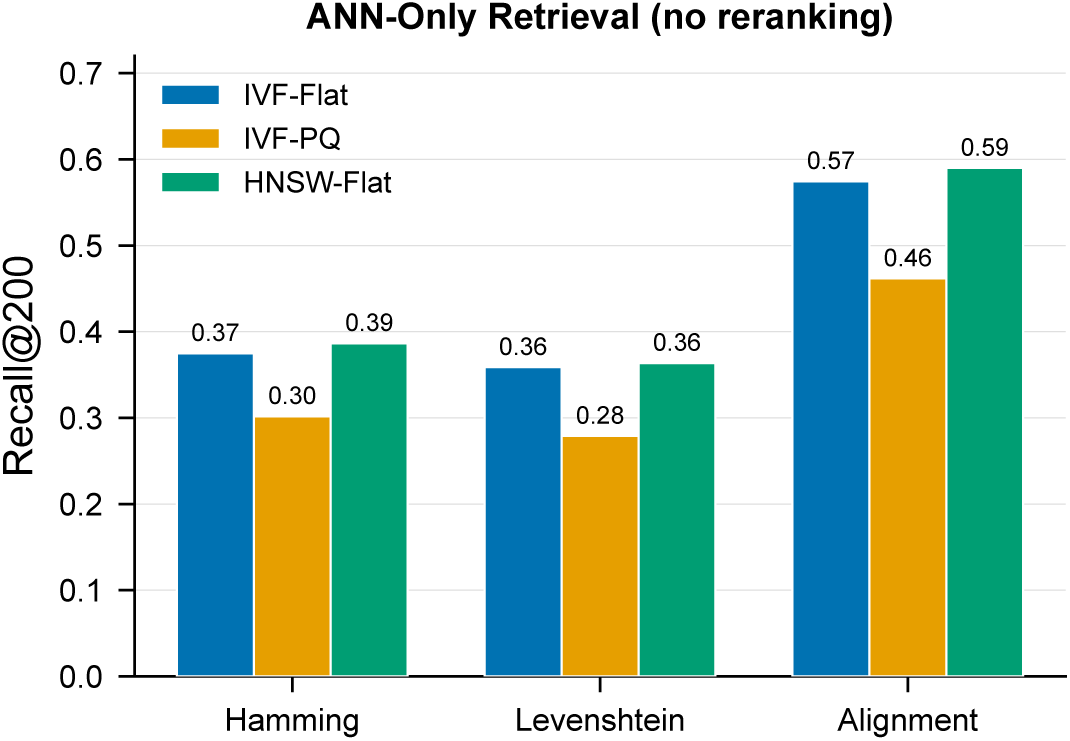
ANN-only retrieval recall@200 across index types and ground truth metrics. The alignment metric yields the highest recall across all index types, reflecting the BLOSUM62-derived embedding’s natural affinity for alignment-based similarity. HNSW-Flat achieves the highest recall, while IVF-PQ offers the lowest latency.

**Table 1:** ANN-only retrieval performance at best search parameter per index.

| Index | Metric | Recall@200 | Median ms | P95 ms |
| --- | --- | --- | --- | --- |
| IVF-Flat | Hamming | 0.3749 | 0.932 | 1.147 |
| IVF-Flat | Levenshtein | 0.3588 | 0.932 | 1.147 |
| IVF-Flat | Alignment | 0.5746 | 0.932 | 1.147 |
| IVF-PQ | Hamming | 0.3018 | 0.032 | 0.052 |
| IVF-PQ | Levenshtein | 0.2790 | 0.032 | 0.052 |
| IVF-PQ | Alignment | 0.4618 | 0.032 | 0.052 |
| HNSW-Flat | Hamming | 0.3865 | 0.738 | 0.827 |
| HNSW-Flat | Levenshtein | 0.3636 | 0.738 | 0.827 |
| HNSW-Flat | Alignment | 0.5900 | 0.738 | 0.827 |

The alignment metric consistently yielded the highest ANN-only recall across all index types, indicating that the BLOSUM62-kernelized embedding is most naturally aligned with alignment-based similarity.

### 2.4 Reranking Dramatically Improves Retrieval Quality

The second-stage exact reranking produced substantial and consistent improvements in exact-top10 recall, particularly at strict top-*k* cutoffs (Table 2, Fig. 3). At *k* = 10, reranking lifted exact-top10 recall by +0.31 to +0.63 absolute across all index-metric combinations. The largest gains were observed for IVF-PQ with Hamming reranking (+0.6310), reflecting the ability of exact reranking to correct ranking errors introduced by product quantization. The highest absolute exact-top10 recall was achieved by HNSW-Flat with alignment reranking (0.9799).

**Figure 3:**
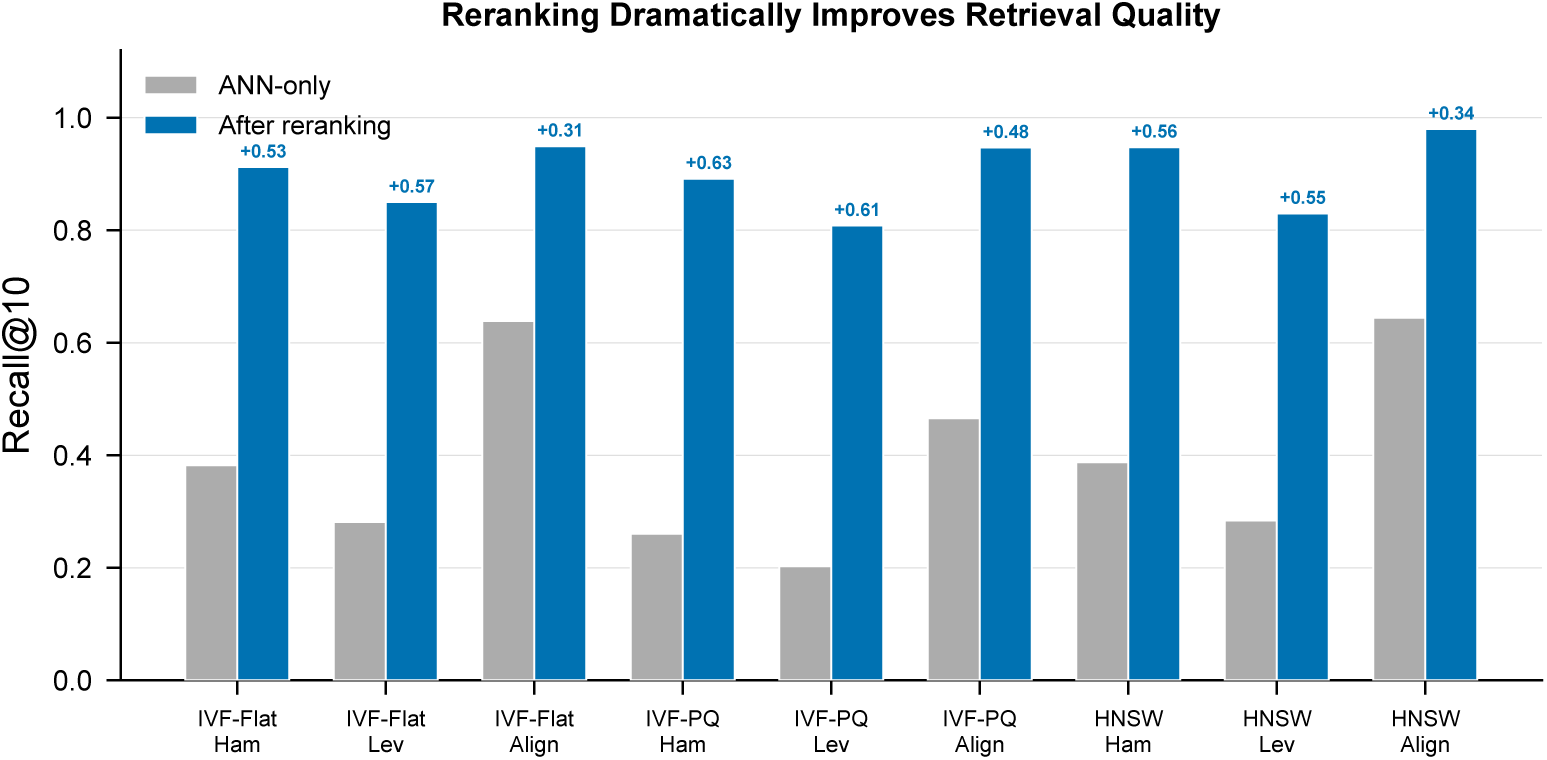
Reranking dramatically improves retrieval quality. Paired comparison of ANN-only exact-top10 recall (grey) versus reranked exact-top10 recall (blue) across all index–metric combinations. Exact-top10 recall is the fraction of the ground-truth top-10 neighbors recovered in the returned top-10. Reranking provides +0.31 to +0.63 absolute recall improvement, with the largest gains for IVF-PQ where product quantization introduces the most ranking error.

**Table 2:** Reranking gains at *k* = 10 with shortlist k = 1,000. Here R@10 denotes exact-top10 recall, i.e. the fraction of the ground-truth top-10 neighbors recovered in the returned top-10.

| Index | Metric | ANN R@10 | Rerank R@10 | Lift | E2E ms |
| --- | --- | --- | --- | --- | --- |
| IVF-Flat | Hamming | 0.3825 | 0.9124 | +0.530 | 1.57 |
| IVF-Flat | Levenshtein | 0.2814 | 0.8501 | +0.569 | 1.66 |
| IVF-Flat | Alignment | 0.6386 | 0.9494 | +0.311 | 4.84 |
| IVF-PQ | Hamming | 0.2607 | 0.8916 | +0.631 | 0.62 |
| IVF-PQ | Levenshtein | 0.2026 | 0.8083 | +0.606 | 0.72 |
| IVF-PQ | Alignment | 0.4657 | 0.9470 | +0.481 | 3.84 |
| HNSW-Flat | Hamming | 0.3877 | 0.9477 | +0.560 | 1.09 |
| HNSW-Flat | Levenshtein | 0.2839 | 0.8299 | +0.546 | 1.20 |
| HNSW-Flat | Alignment | 0.6445 | 0.9799 | +0.335 | 4.50 |

At *k* = 200, reranking continued to improve recall but with diminishing marginal gains (Table 3). HNSW-Flat with alignment reranking achieved recall@200 of 0.8199, compared to 0.5863 for ANN-only (+0.2336). This diminishing-returns pattern is expected: at large *k*, the reranked list spans a wider distance range where the shortlist coverage becomes the binding constraint.

**Table 3:** Reranking performance at *k* = 200.

| Index | Metric | ANN R@200 | Rerank R@200 | Lift | E2E ms |
| --- | --- | --- | --- | --- | --- |
| IVF-Flat | Hamming | 0.3744 | 0.6987 | +0.324 | 1.57 |
| IVF-Flat | Levenshtein | 0.3592 | 0.6557 | +0.297 | 1.66 |
| IVF-Flat | Alignment | 0.5752 | 0.8438 | +0.269 | 4.84 |
| IVF-PQ | Hamming | 0.3006 | 0.6211 | +0.321 | 0.62 |
| IVF-PQ | Levenshtein | 0.2791 | 0.5724 | +0.293 | 0.72 |
| IVF-PQ | Alignment | 0.4607 | 0.7926 | +0.332 | 3.84 |
| HNSW-Flat | Hamming | 0.3844 | 0.6733 | +0.289 | 1.09 |
| HNSW-Flat | Levenshtein | 0.3585 | 0.5803 | +0.222 | 1.20 |
| HNSW-Flat | Alignment | 0.5863 | 0.8199 | +0.234 | 4.50 |

### 2.5 Speedup over Exact Search

To quantify the practical efficiency advantage, we compared TCRseek’s end-to-end latency against exact brute-force computation of each distance metric (Table 4, Fig. 6). TCRseek achieved online speedups ranging from 3.61× (IVF-Flat, Hamming) to 39.58× (IVF-PQ, alignment). The speedup advantage increased with the computational cost of the target metric: alignment-based retrieval, the most expensive exact operation, benefited most from the twostage decomposition, with all three index types achieving *>* 30× speedup.

**Table 4:**
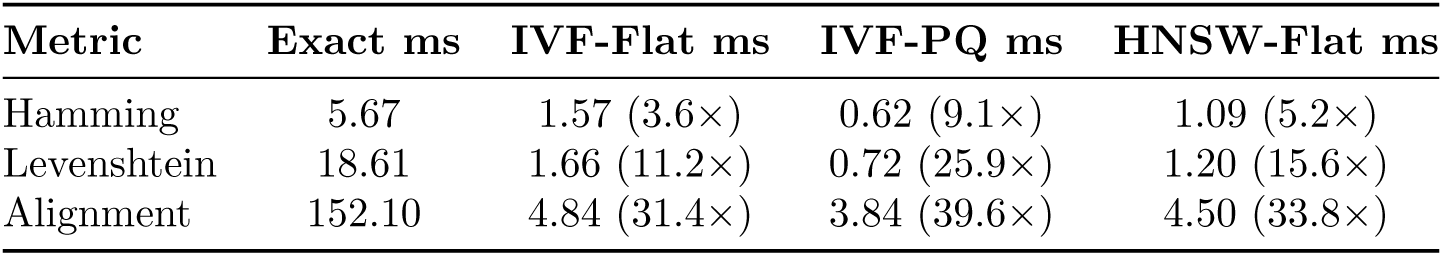
Online speedup of TCRseek (ANN + rerank) versus exact search.

### 2.6 Index Selection Tradeoffs

The three FAISS index architectures offered distinct operating points. IVF-PQ provided the fastest end-to-end throughput (0.62–3.84 ms/query), making it optimal for latency-critical applications. HNSW-Flat achieved the highest ANN-only recall without reranking. IVF-Flat provided a balanced middle ground. For top-10 alignment-distance neighbor retrieval, HNSW-Flat with alignment reranking achieved near-ceiling recall of 0.9799 at 4.50 ms/query, while IVF-PQ offered 0.9470 recall at 3.84 ms/query—only a 3.3% recall reduction for a 15% latency improvement.

### 2.7 Sensitivity of Retrieval Accuracy to Shortlist Size

A key design parameter in TCRseek’s two-stage pipeline is the shortlist size—the number of ANN candidates passed to the exact reranking stage. We systematically varied the shortlist size from 10 to 1,000 and measured two complementary metrics across all three index types and ground truth distances (Fig. 8): binary top-200 recall@10 (top row) and exact-top10 recall (bottom row). Binary top-200 recall@10 is defined as the number of ground-truth top-200 neighbors recovered in the returned top-10 divided by 200, so its theoretical maximum is 10*/*200 = 0.05; it is therefore distinct from the exact-top10 recall values reported in Table 2 and Fig. 3, which quantify recovery of the true top-10 neighbors.

Binary top-200 recall@10 approached its 0.05 ceiling rapidly across all configurations. Under alignment ground truth, the metric rose from 0.0444–0.0478 at shortlist size 10 to 0.0495– 0.0498 at shortlist 20, and reached 0.0500 by shortlist 50–200 depending on backbone. Under Levenshtein ground truth, it approached the same ceiling by shortlist 50 for all three backbones. Hamming distance required larger shortlists, reaching 0.0496–0.0497 by shortlist 100 and 0.0500 only at shortlist 500–1,000.

Exact-top10 recall proved more informative for distinguishing backbone performance. Unlike binary top-200 recall@10, this metric continued to improve with increasing shortlist size, reflecting the progressive inclusion of true nearest neighbors from more distant regions of the embedding space. For example, under alignment ground truth, IVF-PQ increased from 0.631 at shortlist 20 to 0.947 at shortlist 1,000 even though binary top-200 recall@10 changed only from 0.04955 to 0.05000. At the default shortlist size of 1,000, exact-top10 recall under alignment ground truth reached 0.980 (HNSW-Flat), 0.949 (IVF-Flat), and 0.947 (IVF-PQ). Under Hamming ground truth, the values were 0.948 (HNSW-Flat), 0.912 (IVF-Flat), and 0.892 (IVF-PQ), showing clearer separation between index types. These diminishing returns suggest that shortlist sizes of 20–50 are sufficient when the objective is simply to place members of the top-200 relevant set into the returned top-10, whereas larger shortlists (200–500) are warranted when high recovery of the exact top-10 neighbors is important.

### 2.8 Multi-Method Comparison on Exact Ground Truth

To contextualize TCRseek’s retrieval quality relative to existing tools, we conducted a head-to-head multi-method benchmark (Table 5, Fig. 4) comparing TCRseek (IVF-Flat with NW alignment reranking), tcrdist3, TCRMatch, and GIANA. Each method retrieved its top-10 nearest neighbors for 10,000 query sequences from the 100,000-sequence corpus.

**Figure 4:**
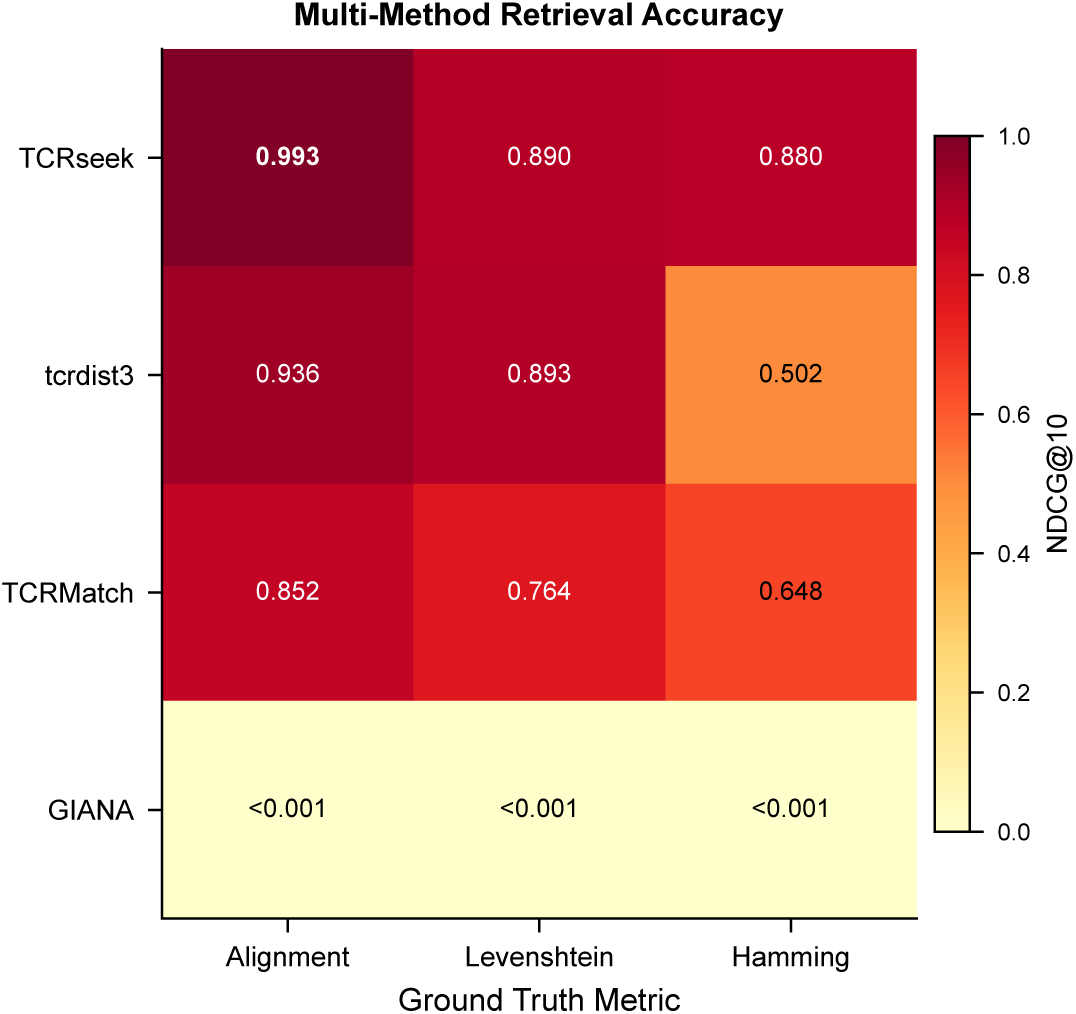
Multi-method retrieval accuracy heatmap. NDCG@10 for TCRseek, tcrdist3, TCR-Match, and GIANA (rows) evaluated against three ground truth distance metrics (columns). TCRseek is strongest under alignment and Hamming ground truth, while TCRseek and tcrdist3 are nearly tied under Levenshtein ground truth.

**Table 5:** Multi-method retrieval accuracy evaluated against exact ground truth under three distance metrics (10,000 queries, top-10 retrieval from 100,000 sequences). TCRseek uses alignment reranking, while tcrdist3, TCRMatch, and GIANA provide sequence-distance or clustering baselines for comparison.

| Method | Ground Truth | P@1 | P@5 | P@10 | MRR | NDCG@10 |
| --- | --- | --- | --- | --- | --- | --- |
| **TCRseek** | Alignment | **0.999** | **0.996** | **0.991** | **0.999** | **0.993** |
| tcrdist3 | Alignment | 0.980 | 0.951 | 0.923 | 0.990 | 0.936 |
| TCRMatch | Alignment | 0.917 | 0.874 | 0.833 | 0.919 | 0.852 |
| GIANA | Alignment | 0.001 | <0.001 | <0.001 | 0.001 | <0.001 |
| tcrdist3 | Levenshtein | 0.953 | 0.910 | **0.877** | 0.975 | **0.894** |
| **TCRseek** | Levenshtein | **0.967** | **0.918** | 0.867 | **0.982** | 0.890 |
| TCRMatch | Levenshtein | 0.884 | 0.800 | 0.730 | 0.900 | 0.764 |
| GIANA | Levenshtein | 0.001 | <0.001 | <0.001 | 0.001 | <0.001 |
| **TCRseek** | Hamming | **0.940** | **0.896** | **0.863** | **0.966** | **0.880** |
| TCRMatch | Hamming | 0.769 | 0.679 | 0.616 | 0.824 | 0.648 |
| tcrdist3 | Hamming | 0.579 | 0.515 | 0.484 | 0.724 | 0.502 |
| GIANA | Hamming | 0.001 | <0.001 | <0.001 | 0.001 | <0.001 |

#### Alignment ground truth (matched-metric)

Under alignment ground truth, TCRseek’s NW reranker uses the same scoring function that defines the ground truth (see Section 4.6) and achieved the strongest retrieval performance among the retained baselines (P@10 = 0.991, MRR = 0.999, NDCG@10 = 0.993). tcrdist3 ranked second (NDCG@10 = 0.936), followed by TCRMatch (0.852). GIANA, a clustering-oriented method not designed for ranked nearest-neighbor retrieval under these metrics, returned near-zero precision (P@10 *<* 0.001; see Section 2.10).

#### Levenshtein ground truth (cross-metric)

Under Levenshtein ground truth—where TCRseek’s alignment-based reranker differs from the evaluation metric—tcrdist3 (NDCG@10 = 0.894) and TCRseek (0.890) were nearly indistinguishable, indicating similar generalization to edit-distance-based similarity. TCRMatch dropped to 0.764, while GIANA again remained near zero.

#### Hamming ground truth (cross-metric)

Under Hamming ground truth, TCRseek ranked highest (NDCG@10 = 0.880), followed by TCRMatch (0.648), tcrdist3 (0.502), and GIANA near zero. This ordering suggests that the BLOSUM62-derived embedding captures positional mismatch patterns that correlate well with Hamming distance and substantially better preserves Hamming-style neighborhood structure than the alternative retained baselines.

### 2.9 Precision Degradation with Increasing k

The multi-method comparison revealed characteristic precision decay curves that differ markedly across methods (Table 6, Fig. 5). Under alignment ground truth, TCRseek maintained P@*k* above 0.90 through *k* = 80, declining to 0.88 at *k* = 100 and 0.69 at *k* = 200. tcrdist3 decayed more gradually at early *k* but remained below TCRseek throughout alignment-groundtruth retrieval, while TCRMatch dropped more steeply from P@10 = 0.833 to P@100 = 0.502 and P@200 = 0.354. Under Hamming ground truth, TCRMatch remained consistently above tcrdist3 but well below TCRseek. For applications requiring only a small number of nearest neighbors (*k* `≤` 20), TCRseek provides near-perfect retrieval under alignment distance and remains the strongest retained retrieval method.

**Figure 5:**
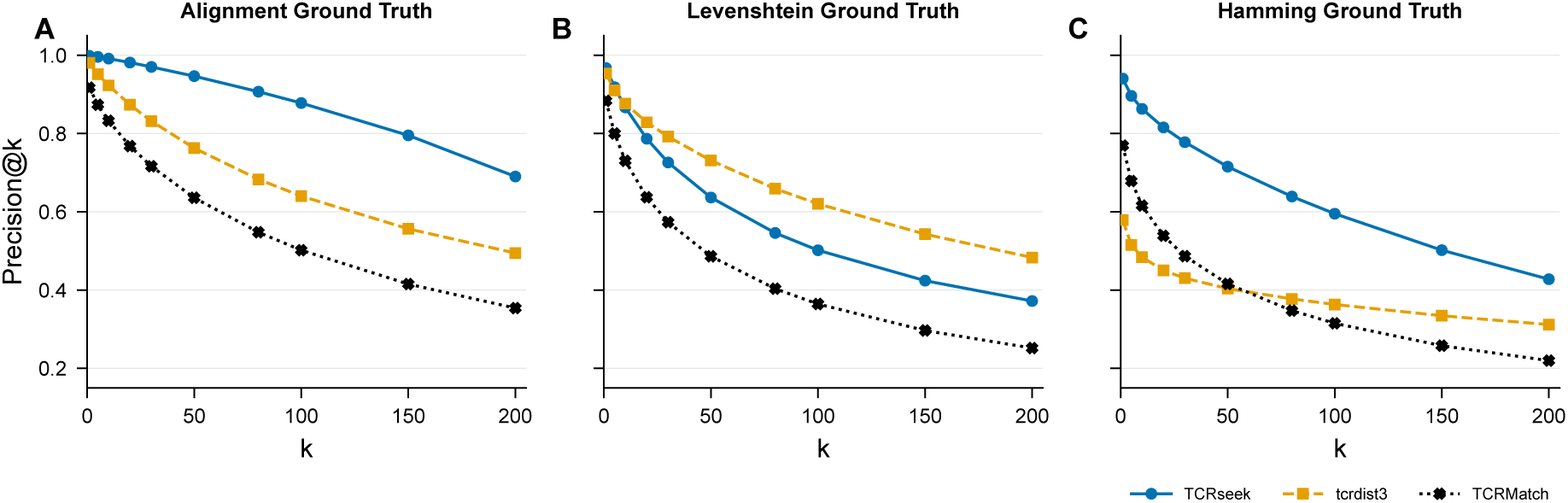
Precision@*k* curves across methods and ground truth metrics. (A) Alignment, (B) Levenshtein, and (C) Hamming ground truth. TCRseek (blue) remains strongest under alignment and Hamming ground truth, while TCRseek and tcrdist3 are closely matched under Levenshtein ground truth. GIANA is omitted because its precision remained near zero across all *k* values (see Section 2.10).

**Figure 6:**
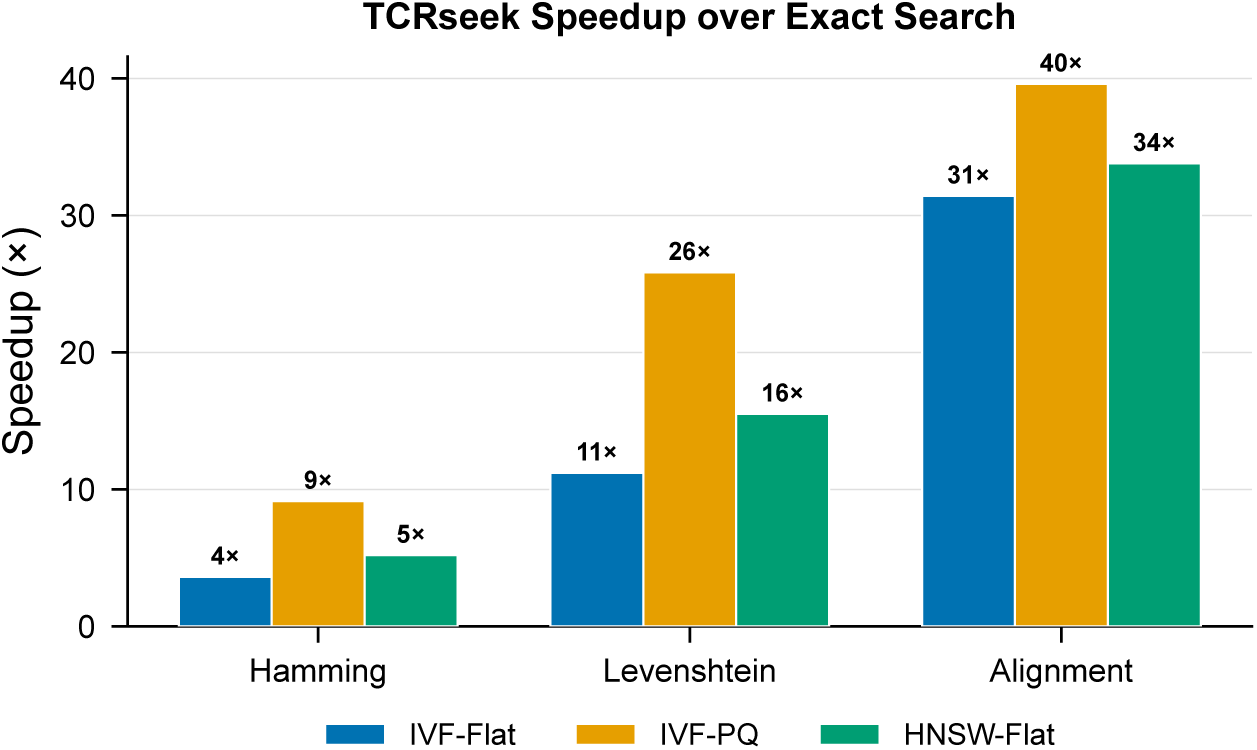
TCRseek speedup over exact search by index type and distance metric. Speedup increases with the computational cost of the target metric, reaching up to 40× for alignment distance with IVF-PQ.

**Figure 7:**
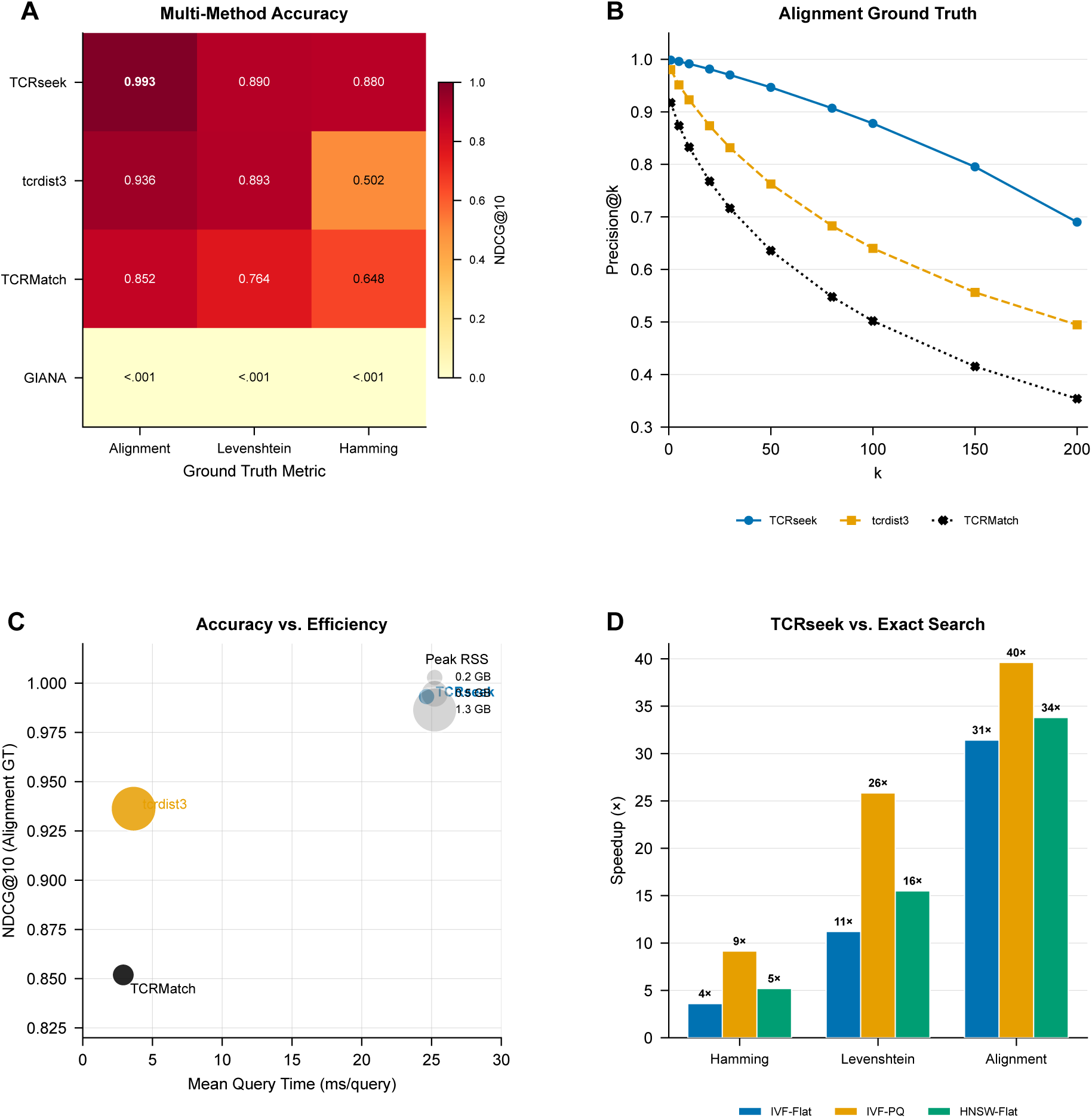
Combined results overview. (A) Multi-method NDCG@10 heatmap across three ground truth metrics. (B) Precision@*k* curves under alignment ground truth. (C) Accuracy-efficiency comparison on the shared stage-A benchmark. (D) TCRseek speedup over exact search by index type and distance metric.

**Figure 8:**
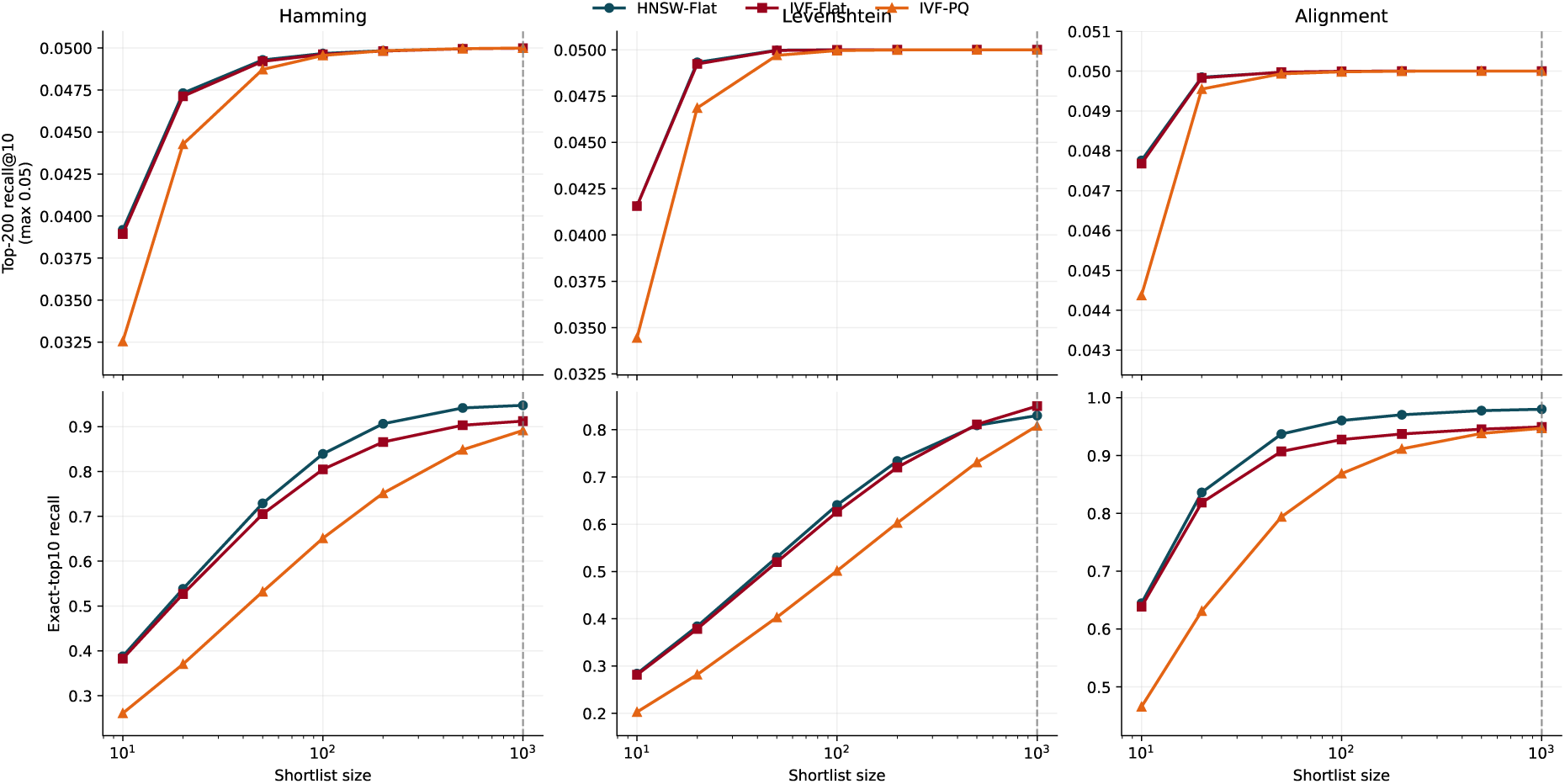
Sensitivity of retrieval accuracy to shortlist size. Top row: binary top-200 recall@10, defined as the number of ground-truth top-200 neighbors recovered in the returned top-10 divided by 200; its theoretical maximum is 0.05. Bottom row: exact-top10 recall, defined as the fraction of true top-10 neighbors recovered in the reranked output. Columns correspond to Hamming, Levenshtein, and alignment ground truth metrics. Binary top-200 recall@10 approaches its 0.05 ceiling rapidly, whereas exact-top10 recall continues to improve with larger shortlists and more clearly separates index types.

**Table 6:** Precision@ *k* across methods for alignment and Hamming ground truth at selected *k* values. GIANA is omitted because its precision remained near zero across all *k* values.

| Method | Ground Truth | P@1 | P@10 | P@50 | P@100 | P@200 |
| --- | --- | --- | --- | --- | --- | --- |
| **TCRseek** | Alignment | **0.999** | **0.991** | **0.947** | **0.878** | **0.690** |
| tcrdist3 | Alignment | 0.980 | 0.923 | 0.763 | 0.640 | 0.494 |
| TCRMatch | Alignment | 0.917 | 0.833 | 0.636 | 0.502 | 0.354 |
| **TCRseek** | Hamming | **0.940** | **0.863** | **0.715** | **0.595** | **0.428** |
| TCRMatch | Hamming | 0.769 | 0.616 | 0.416 | 0.315 | 0.220 |
| tcrdist3 | Hamming | 0.579 | 0.484 | 0.404 | 0.363 | 0.312 |

### 2.10 GIANA: Distance-Metric Mismatch

GIANA exhibited near-zero precision across all three ground truth metrics (P@10 *<* 0.002). This result should be interpreted with caution: GIANA was designed as a clustering tool for identifying groups of functionally related TCRs via isometric encoding, not as a ranked nearest-neighbor retrieval system optimized for Hamming, Levenshtein, or alignment distances. The near-zero scores therefore reflect a mismatch between GIANA’s design objective and the retrieval-oriented evaluation framework used here, rather than a deficiency in GIANA’s intended use case. We include GIANA in the comparison to illustrate that methods designed for different tasks (clustering vs. ranked retrieval) are not interchangeable when evaluated under explicit distance-based ground truth.

## 3 Discussion

We have presented TCRseek, a framework for scalable TCR repertoire analysis that combines multi-scale windowed k-mer embedding with FAISS-based approximate nearest neighbor search and exact sequence-level reranking. The two-stage architecture addresses a fundamental tension in computational immunology: the need for biologically meaningful sequence comparison (which requires expensive alignment computation) and the need for practical scalability (which demands sublinear search algorithms). By separating these concerns—using fast but approximate embedding-space search for candidate generation, followed by rigorous but expensive alignment-based reranking—TCRseek achieves both goals simultaneously.

An important nuance of the benchmark design is that TCRseek’s alignment-based reranker uses the same Needleman–Wunsch scoring function that defines the alignment ground truth. In this matched-metric setting, TCRseek’s NDCG@10 of 0.993 indicates that the remaining error arises primarily from true neighbors missed by the ANN shortlist rather than from reranking inaccuracy. The more informative evaluation is the cross-metric setting, where the reranking metric differs from the ground truth. Under Levenshtein ground truth, TCRseek (NDCG@10 = 0.890) performed comparably to tcrdist3 (0.894) and substantially better than TCRMatch (0.764), while under Hamming ground truth, TCRseek (0.880) substantially outperformed TCRMatch (0.648) and tcrdist3 (0.502). These cross-metric results demonstrate that TCRseek’s BLOSUM62-derived embedding captures sequence similarity patterns that generalize beyond the specific distance metric used for reranking. No single retained method dominated across all distance definitions, consistent with the expectation that different distance metrics capture distinct aspects of sequence similarity.

The choice of BLOSUM62 eigendecomposition as the basis for amino acid representation is deliberate. Unlike arbitrary embeddings or learned representations, BLOSUM62 vectors encode substitution patterns observed across conserved protein blocks, making them directly relevant to the biochemistry of TCR-pMHC recognition. Projecting the centered BLOSUM62 matrix onto its positive eigenspace yields Euclidean coordinates that approximate centered BLOSUM62 similarities in a principled manner, without the need for training data or model fitting.

The multi-scale windowed k-mer design addresses the variable-length nature of CDR3 sequences. By assigning k-mers to positional windows, the embedding captures not only which amino acid motifs are present (compositional information) but also where they occur along the CDR3 loop (positional information). This architecture is related to spatial pyramid matching in computer vision (Lazebnik et al., 2006), adapted here for protein sequence analysis.

The reranking stage is essential for biological accuracy. Our results demonstrated that embedding-space nearest neighbors alone provide moderate recall of true alignment-space neighbors, but exact reranking corrects ranking errors. The cost of reranking a shortlist of 200 candidates is modest compared to the savings from avoiding exhaustive pairwise comparison over the full database.

An important architectural advantage of TCRseek is its sublinear query-time scaling. Methods based on exact pairwise computation—including tcrdist3 and the brute-force baselines— exhibit *O*(*N*) per-query scaling, which becomes prohibitive as corpus sizes grow. TCRseek’s ANN index ensures that query latency grows sublinearly with database size, and the speedup analysis (Table 4) demonstrated 3.6–39.6× speedups over exact search even at the 100,000-sequence scale, with the largest gains for alignment-based retrieval. GIANA’s near-zero retrieval precision illustrates that methods designed for clustering are not interchangeable with ranked nearest-neighbor retrieval systems, even when both operate on CDR3 sequence similarity.

Several limitations should be acknowledged. First, TCRseek currently operates on CDR3*β* sequences alone; incorporating paired alpha-beta chain information could improve specificity. Second, the embedding parameters (*k* ∈ {3, 4, 5}, *B* ∈ {3, 5, 10}, *d* = 19) were selected based on domain knowledge rather than optimized on benchmark data; a systematic ablation study varying these parameters would clarify the contribution of each multi-scale branch and determine whether a lower-dimensional embedding could achieve comparable retrieval quality at reduced computational cost. Third, the 4,104-dimensional embedding is high-dimensional relative to typical ANN workloads, and the modest ANN-only recall@200 (0.37–0.59) suggests that dimensionality reduction (e.g., PCA to 128–512 dimensions) warrants investigation. Fourth, all benchmark metrics are reported as point estimates without confidence intervals; bootstrap resampling or repeated random query-set selection would quantify the statistical significance of inter-method differences, particularly where methods appear close (e.g., TCRseek vs. tcrdist3 under Levenshtein ground truth). Fifth, the current implementation uses CPU-only FAISS; GPU acceleration would further improve throughput. Finally, although the benchmark now includes TCRMatch as a sequence-kernel baseline, it still does not include several other ANN-adapted TCR methods or learned retrieval backbones beyond the sequence-only baselines considered here, and the comparison with GIANA—a clustering tool not designed for ranked retrieval—should be interpreted in light of this design mismatch.

Future work will extend TCRseek to paired-chain TCR data, conduct systematic ablation studies over embedding parameters and dimensionality reduction, benchmark against additional deep learning methods (TCR-BERT, DeepTCR, TouCAN) and other ANN-adapted TCR tools beyond TCRMatch, report bootstrap confidence intervals for all retrieval metrics, explore learned embeddings as alternatives to the BLOSUM62-based scheme, and develop a streaming mode for real-time repertoire monitoring.

In conclusion, TCRseek demonstrates that approximate nearest neighbor search, when combined with biologically informed embeddings and exact reranking, provides a practical and scalable framework for TCR repertoire analysis. The two-stage design achieves near-perfect retrieval when the reranking metric matches the evaluation criterion, and competitive cross-metric generalization otherwise, with substantial speedups over exact brute-force search. By making million-sequence nearest-neighbor search accessible on standard hardware, this approach opens the door to population-scale studies of T-cell immunity that were previously computationally intractable.

## 4 Methods

### 4.1 Amino Acid Embedding via BLOSUM62 Eigendecomposition

The foundation of the TCRseek embedding is a biologically informed numerical representation of individual amino acid residues. Rather than using arbitrary one-hot encodings, which discard information about physicochemical similarity between residues, we derive amino acid vectors from the BLOSUM62 substitution matrix (Henikoff & Henikoff, 1992), a standard scoring matrix that captures evolutionary substitution patterns observed in conserved protein blocks.

Let **S** denote the 20 × 20 BLOSUM62 matrix indexed by the 20 canonical amino acids. We first apply centering to form **K** = **HSH**, where **H** = **I** − (1*/*20)**11**^⊤^ is the centering matrix. For the bundled BLOSUM62 matrix used in this work, the centered matrix has 19 positive eigenvalues and one numerical zero eigenvalue at the implementation tolerance. We then compute the eigendecomposition **K** = **VΛV**^⊤^ and retain only the *d* eigenvectors corresponding to the positive eigenvalues, scaled by the square root of their eigenvalues: each amino acid *a* is represented by the *d*-dimensional row vector 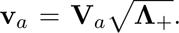 In the default configuration, we retain *d* = 19 dimensions, so the full positive spectrum is kept and the zero mode is discarded. The resulting positive-spectrum embedding yields inner products ⟨**v***_a_,* **v***_b_*⟩ that approximate the centered BLOSUM62 similarity between amino acids *a* and *b*, providing an embedding where Euclidean distances reflect substitution penalties directly relevant to protein function.

### 4.2 Multi-Scale Windowed k-mer Embedding

Given a CDR3 amino acid sequence *s* = *s*_1_*s*_2_ *. . . s_L_*of length *L*, we construct a fixed-length embedding through a multi-scale windowed k-mer aggregation scheme. For each combination of k-mer size *k* ∈ {3, 4, 5} and window count *B* ∈ {3, 5, 10}, we define *B* positional windows that partition the CDR3 sequence into contiguous regions. Each k-mer starting at position *i* is assigned to window *w* = ⌊(*i/* max(1*, L* − *k*)) · *B*⌋, clipped to [0*, B* − 1]. Optionally, soft-linear window assignment distributes each k-mer across adjacent windows using distance-weighted contributions, providing smoother positional representation.

For each k-mer *m* = *s_i_s_i_*_+1_ *. . . s_i_*_+*k*−1_, we form the concatenated amino acid vector **x***_m_* = [**v***_s_* ; **v***_s_* ; *. . .* ; **v***_s_*] ∈ R*^kd^*. K-mer vectors within each window are aggregated by summation, and each window vector is L2-normalized before concatenation. The concatenated vector across all *B* windows for a single (*k, B*) branch has dimension *B* ·*k* ·*d*. The full embedding concatenates all branches across the multi-scale grid of (*k, B*) values, yielding a vector of dimension ∑*_k,B_ B* · *k* · *d*. With the default parameters (*k* ∈ {3, 4, 5}, *B* ∈ {3, 5, 10}, *d* = 19), this produces a vector of 4,104 dimensions. A final global L2-normalization maps the concatenated vector to the unit hypersphere, enabling cosine-like similarity search via Euclidean distance.

This multi-scale design captures sequence features at different resolutions: small *k* values detect local residue patterns while large *k* values capture longer-range motifs; few windows (*B* = 3) provide coarse positional information while many windows (*B* = 10) distinguish finegrained CDR3 subregions such as the conserved N-terminal cysteine region, the hypervariable central loop, and the conserved C-terminal phenylalanine-glycine motif.

### 4.3 FAISS Index Construction

The embedded CDR3 vectors are indexed using the FAISS library (Johnson et al., 2019) for approximate nearest neighbor search. TCRseek supports three index architectures:

#### IVF-Flat

An inverted file index partitions the embedding space into *nlist* Voronoi cells using k-means clustering, then stores vectors within their assigned cells. At query time, only the *nprobe* closest cells are searched exhaustively, reducing the search space from *N* to approximately *N* · nprobe*/*nlist vectors. With default parameters nlist = 256 and nprobe = 16, this provides a 16× reduction in comparisons while maintaining high recall.

#### IVF-PQ

Inverted file indexing is combined with product quantization, which compresses each vector into *m* sub-quantizer codes of *nbits* bits each. This reduces both memory footprint and search cost by replacing exact distance computation with table lookups on quantized subvectors. For embeddings not evenly divisible by *m*, the vectors are right-padded with zeros to the nearest multiple. IVF-PQ is particularly beneficial for large-scale deployments where memory constraints preclude storing full-precision vectors.

#### HNSW-Flat

A hierarchical navigable small world graph (Malkov & Yashunin, 2020) constructs a multi-layer proximity graph over the full-precision vectors. At construction time, each vector is inserted into the graph with *M* bidirectional connections per layer (default *M* = 32) and a construction-time search width of efConstruction = 200. At query time, the graph is traversed with a beam width of efSearch (evaluated at 32, 64, and 128). HNSW-Flat achieves higher recall than IVF-based indices at comparable latency, at the cost of a larger memory footprint and longer index construction time.

Index construction proceeds deterministically: training vectors for centroid computation (IVF-based indices) and graph construction (HNSW) are drawn using a fixed random seed, ensuring reproducible index builds. The index, along with associated metadata (sequence IDs, raw sequences, embedding configuration), is persisted to a structured artifact directory for reuse.

### 4.4 Two-Stage Retrieval with Reranking

For each query CDR3 sequence, the retrieval process proceeds in two stages:

#### Stage 1: ANN shortlisting

The query sequence is embedded using the same multiscale windowed k-mer procedure applied to the database, and the FAISS index returns the top *shortlist k* approximate nearest neighbors (default shortlist k = 200). Self-matches (queries appearing in the database) are excluded.

#### Stage 2: Exact reranking

The shortlisted candidates are rescored using an exact sequence-level distance metric. TCRseek supports four reranking metrics:

- **Needleman–Wunsch (NW) alignment**: Global alignment using the BLOSUM62 substitution matrix with affine gap penalties (default gap open = −10, gap extend = −1), optionally accelerated by banded alignment. The alignment score is negated to convert similarity to distance. This is the default reranking metric and provides the most biologically meaningful ranking.
- **Smith–Waterman (SW) alignment**: Local alignment with the same parameters, capturing partial sequence similarity.
- **Levenshtein distance**: Standard edit distance counting insertions, deletions, and substitutions.
- **Hamming distance**: Positional mismatch count for same-length sequences, with length-penalized extension for sequences of different lengths.

Before reranking, length-based prefiltering removes candidates whose length differs from the query by more than a configurable threshold, avoiding unnecessary computation on clearly dissimilar pairs. The final output is a ranked list of the top *k* neighbors per query, ordered by reranking distance.

### 4.5 Benchmark Framework

We developed a comprehensive benchmark framework to evaluate TCRseek against existing TCR analysis methods across two tracks: retrieval accuracy and retrieval efficiency.

#### Datasets

The retrieval accuracy benchmark corpus was derived from a large-scale TCR sequencing dataset. We filtered for human TRB chain sequences with canonical amino acid CDR3 regions and applied deterministic hash-based sampling to select 100,000 unique sequences for the ground-truth-locked accuracy evaluation.

#### Compared methods

The benchmark included adapters for the following methods:

- **TCRseek** (this work): IVF-Flat, IVF-PQ, and HNSW-Flat indexing with NW reranking
- **tcrdist3** (Mayer-Blackwell et al., 2021): Sparse rectangular CDR3 distance with BLOSUM62-based alignment
- **TCRMatch** (Chronister et al., 2021): Sequence-similarity kernel for TCR matching
- **GIANA** (Zhang et al., 2021): Isometric CDR3 encoding designed for clustering; included to assess cross-paradigm comparability

#### Retrieval metrics

For each query, we computed precision@*k* (*k* = 1, 5, 10), recall@*k* (*k* = 5, 10), mean reciprocal rank (MRR), and NDCG@10 (normalized discounted cumulative gain).

### 4.6 Computational complexity of benchmarked methods

Mathematically, the main difference among the benchmarked methods is where the computational burden sits. TCRseek, implemented here through the `tcrfaiss` family of backends, pays for a two-stage pipeline consisting of sequence embedding, approximate nearest-neighbor (ANN) shortlist generation, and exact reranking. By contrast, `tcrdist3` uses a cheaper biological pairwise distance than TCRMatch but still incurs near-exhaustive query cost when applied directly against a large reference set. GIANA attempts to move the problem into a transformed Euclidean space so that search can be handled geometrically rather than by direct biological comparison, whereas TCRMatch remains dominated by an intrinsically expensive sequence kernel. Exact Hamming and Levenshtein calculations remain useful reference computations for ground-truth construction and speedup analysis, but their exhaustive formulations do not scale favorably.

For TCRseek, the end-to-end query cost can be written as

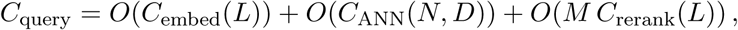

where *N* is the reference size, *L* is the typical CDR3 length, *D* is the fixed embedding dimension, and *M* is the shortlist size returned by the ANN stage. With fixed branch sets and a fixed amino-acid embedding dimension, the windowed *k*-mer encoder is linear in sequence length,

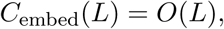

because it scans each valid *k*-mer only once per configured branch. The dominant remaining term is therefore determined by the index backend and the exact reranker. In the benchmark configurations, reranking is typically performed with Needleman–Wunsch alignment, so *C*_rerank_(*L*) = *O*(*L*^2^). Consequently, the practical per-query cost of TCRseek is usually best understood as

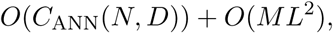

rather than as ANN search alone.

Within the `tcrfaiss` family, the precise ANN term depends on the backend. For IVF-Flat, a useful approximation is

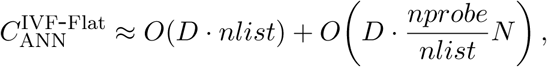

reflecting coarse-centroid assignment followed by scanning the probed inverted lists. For IVFPQ, the coarse stage is similar, but list scanning is performed on compressed product-quantized codes, giving a query cost of approximately

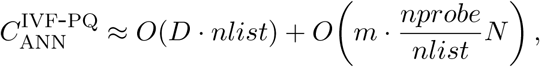

where *m* is the number of subquantizers. HNSW-Flat has expected sublinear, approximately logarithmic behavior in *N* , often summarized as *O*(*D ef Search* log *N*) per query, with a higher graph-construction cost during indexing. The FAISS LSH backend is the important exception: because `IndexLSH` is a flat binary-code index rather than a learned partitioning structure, it is not truly sublinear in this benchmark. Its query time remains effectively exhaustive in *N* , albeit with cheaper binary comparisons than floating-point distance evaluation.

The `tcrdist3` baseline occupies an intermediate position. In this benchmark, the adapter computes sparse rectangular CDR3-only distances between all *Q* queries and all *N* references and then extracts the top hits from each sparse row. Its worst-case time complexity is therefore

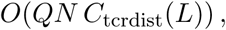

where *C*_tcrdist_(*L*) is the cost of one CDR3 distance computation. For the CDR3-only setting used here, this per-pair cost is much smaller than TCRMatch and is typically close to linear in *L* in practice, although naive optimization over gap placement can make the worst case appear closer to quadratic. Because the implementation stores only neighbors within a radius threshold, the realized memory and postprocessing costs are output-sensitive,

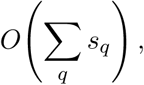

where *s_q_* is the number of retained neighbors for query *q*. Nevertheless, if the radius is permissive or the repertoire is dense, the method still trends toward exhaustive linear-in-*N* search per query and quadratic dataset-level work for neighborhood construction.

GIANA changes the geometry of the problem by encoding each CDR3 into a transformed Euclidean representation. If *r* denotes the transformed-space dimension, then encoding one sequence is approximately *O*(*Lr*), and indexed nearest-neighbor search is expected to scale roughly as *O*(*r* log *N*) rather than *O*(*N*) for each query. At a high level, its cost profile can therefore be summarized as

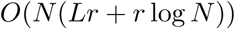

for reference preprocessing and

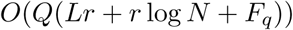

for query-time retrieval, where *F_q_* denotes relatively small cluster or postfilter work. In the present benchmark implementation, retrieval includes both reference clustering and query execution within the same timed run, so the measured GIANA retrieval cost is best interpreted as a combined build-plus-query cost rather than a pure online latency.

TCRMatch remains the most expensive sequence-level comparator among the benchmarked retrieval methods. For two sequences of lengths *m* and *n*, the straightforward kernel computation evaluates all pairs of *k*-mers over all admissible *k*, giving

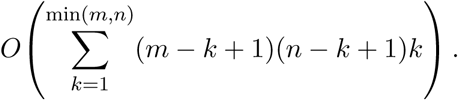

For equal-length sequences of length *L*, this simplifies to

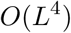

per pair in the straightforward formulation. An exhaustive query batch against a reference database of size *N* therefore scales as

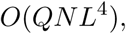

which explains why TCRMatch is best viewed as a high-quality exact scorer for moderate-scale retrieval rather than as a large-scale indexing scheme. Thresholding can reduce the number of reported matches in practice, but it does not change the worst-case asymptotic burden of the underlying kernel comparisons.

Exact Levenshtein and Hamming reference computations are fundamentally exhaustive. Standard Levenshtein distance uses dynamic programming and costs *O*(*L*^2^) per pair, so an all-pairs top-*k* computation scales as

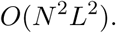

In this benchmark implementation, the Levenshtein adapter computes full reference-to-reference rankings and only then subsets the requested query rows, so its runtime behavior is closer to full *N* ^2^ preprocessing than to a direct *Q* × *N* scan. The Hamming baseline is more favorable only because it buckets sequences by exact length and compares sequences only within the same bucket, yielding

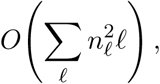

where *n_ℓ_*is the number of sequences of length *ℓ*. In the worst case, this still becomes *O*(*N* ^2^*L*). These reference computations are therefore best interpreted as complexity anchors rather than scalable search methods.

Taken together, the benchmarked methods follow a simple complexity hierarchy. Exact exhaustive approaches, such as TCRMatch, spend nearly all of their effort in direct biological comparison and scale poorly with *N*. `tcrdist3` lowers the per-pair biological cost substantially, but still approaches exhaustive search behavior when used directly against large reference collections. GIANA and the ANN backends of `TCRseek` shift most of the search burden into vector-space indexing, so that only a small candidate set must be evaluated exactly. In this sense, the core asymptotic advantage of the ANN-based design is the replacement of

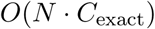

With

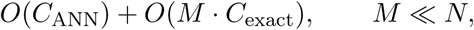

which is precisely the regime required for repertoire-scale TCR retrieval.

### 4.7 Ground Truth Generation

For exact evaluation of retrieval accuracy, we precomputed ground truth top-*k* rankings under multiple distance metrics (Hamming, Levenshtein, and BLOSUM62 global alignment) using a parallelized exact computation pipeline. Alignment distances were defined as *d*(*i, j*) = max(self*_i_,* self*_j_*) − score*_ij_*, clamped at zero, where self*_i_*is the self-alignment score and score*_ij_* is the pairwise Needleman–Wunsch score. This quantity is a symmetric score-derived dissimilarity rather than a canonical metric-space distance. We chose max(self*_i_,* self*_j_*) as the anchor rather than min() or the arithmetic mean because it anchors each pair to the stronger self-alignment baseline, preventing dissimilarity from being diluted when the two sequences have markedly different self-scores due to length or composition differences. In preliminary comparisons, the Spearman rank correlation between max-anchored and mean-anchored rankings exceeded 0.99 across sampled query neighborhoods, indicating that this choice does not materially alter ground truth rankings for the present benchmark. We do not rely on triangle inequality or other metric axioms in the retrieval benchmark; only a consistent ranking function is required. Clamping at zero is included as a numerical safeguard against finite-precision or aligner edge cases; we verified across the full 100,000-sequence benchmark that no pairwise distance was clamped, confirming that the safeguard did not alter any ground truth ranking.

#### Relationship between reranking and ground truth

An important design consideration is that when TCRseek’s Stage 2 reranking uses the Needleman–Wunsch alignment metric, and the ground truth is also defined by the same alignment distance, the reranker has access to the exact scoring function that defines correctness. In this matched-metric setting, the remaining retrieval error arises solely from the ANN shortlist’s failure to include true neighbors, not from reranking inaccuracy. The cross-metric evaluations (Levenshtein and Hamming ground truth) provide the more informative assessment of TCRseek’s generalization ability, because the reranking metric differs from the ground truth definition.

### 4.8 Implementation

TCRseek is implemented as the Python package `TCRseek` with minimal dependencies: NumPy (`≥` 1.24), pandas (`≥` 1.5), FAISS-CPU (1.7.4), Biopython (`≥` 1.81), SciPy (`≥` 1.10), and PyArrow (`≥` 12.0). The benchmark framework additionally requires scikit-learn and psutil. All benchmarks were conducted on a single machine with an Apple M1 Pro processor (10 cores), 16 GB unified memory, running macOS 14.5 and Python 3.11. All code supports Python `≥` 3.9 and is deterministic given fixed random seeds. We note that FAISS k-means initialization is not guaranteed to be bitwise identical across platforms or BLAS implementations; exact numerical reproduction therefore requires matching the FAISS version and linear algebra backend.

## Code and data availability

The TCRseek source code and benchmark framework are available at https://github.com/[repository-url]. The precomputed ground truth files and benchmark configurations are included in the repository to enable independent reproduction of all results reported in this work.

